# A pharmacokinetic-pharmacodynamic assessment of the hepatic and bone-marrow toxicities of the new trypanoside fexinidazole

**DOI:** 10.1101/486761

**Authors:** James A Watson, Nathalie Strub-Wourgraft, Antoine Tarral, Isabela Ribeiro, Joel Tarning, Nicholas J White

## Abstract

Fexinidazole is a novel oral treatment for *Trypanosoma brucei gambiense* human African trypanosomiasis: *g*-HAT. Fexinidazole is also active against other kinetoplastid parasites, notably *T. cruzi* the causative agent of Chagas disease.During the course of a dose ranging assessment in chronic indeterminate Chagas disease, delayed neutropenia and significant increases in hepatic transaminases were observed and clinical investigations were suspended. We retrospectively analyzed all available pharmacokinetic and pharmacodynamic data on fexinidazole in normal healthy volunteers and in patients with chronic Chagas disease and *g*-HAT, in order to assess the determinants of toxicity.

A population pharmacokinetic model was fitted to plasma concentration data on the bioactive fexinidazole sulfone metabolite from three phase 1 studies, two *g*-HAT phase 2/3 field trials and one Chagas phase 2 field trial (462 individuals in total). Bayesian exposure-response models were then fitted to hematological and liver related pharmacodynamic outcomes in chronic Chagas patients.

Neutropenia, reductions in platelet counts, and elevations in liver transaminases were all found to be exposure and thus dose dependent in patients with chronic Chagas disease. Clinically insignificant transient reductions in neutrophil and platelet counts consistent with these exposure-response relationships were observed in the *g*-HAT trials. In contrast, no evidence of hepatotoxicity was observed in the *g*-HAT trials.

Fexinidazole treatment results in a dose dependent liver toxicity and transient bone marrow suppression in Chagas disease. Regimens of shorter duration should be trialled for Chagas. The currently recommended regimen for sleeping sickness provides exposures within a satisfactory safety margin for bone marrow suppression and does not cause hepatotoxicity.

## Introduction

The nitroimidazole fexinidazole is a promising new treatment for sleeping sickness (human African trypanosomiasis caused by *Trypanosoma brucei gambiense*: *g*-HAT). Fexinidazole has the potential to replace current first-line therapies (1; 2; 3) as it is an oral treatment for both the blood stage and the central nervous system (CNS) stages of the disease. Fexinidazole is metabolized extensively *in vivo* to two biologically active metabolites, a sulfoxide (M1) and a more slowly eliminated sulfone (M2) (4; 5). The sulfone metabolite accounts for the majority of bioactive exposure during the ten days of fexinidazole treatment currently recommended in *g*-HAT (6). Extensive phase 2 & 3 studies of fexinidazole in *g*-HAT infections have shown excellent efficacy and good tolerability. This class of drugs also has good *in vitro* and *in vivo* (murine model) activity against other kinetoplastid parasites: *T. cruzi* (5), *T. lewisi* (7) and *Leishmania donovani* (8). Clinical studies have been performed in both Chagas disease and visceral leishmaniasis. In the course of an extended dose finding study in Chagas disease, increases in hepatic transaminases and a delayed and transient fall in neutrophil counts were noted. In response to these findings, clinical studies in Chagas disease were halted temporarily and additional hematology and “liver function” investigations were added to ongoing studies in *g*-HAT. As the prospective studies had not anticipated this toxicity, a retrospective pharmacokinetic-pharmacodynamic assessment was conducted using all the available clinical data to characterize the relationships between drug and metabolite exposure and potential adverse effects, and to provide predictions for the future safety of the ten day fexinidazole regimen developed for the treatment of *g*-HAT.

## Results

### Population pharmacokinetic model of fexinidazole sulfone (M2)

Pharmacokinetic data quantifying concentrations of the metabolite M2 in 462 individuals over more than 4500 time points were analyzed jointly. The overall summary of the six datasets used in this analysis is shown in Table 1.

**Table 1:** Summary of the available pharmacokinetic data. These relate only to the quantification of the metabolite M2. The phase 1 data were used to evaluate the best structural model, and the full dataset was used in the pharmacokinetic analyses in order to estimate the drug exposures in fexinidazole treated chronic Chagas patients and fexinidazole treated *g*-HAT patients.

|  | Trial name | Clinical trial ID | N | Males | Females | Samples per person (median, range) |
| --- | --- | --- | --- | --- | --- | --- |
| Phase 1 | FEX001 | NCT00982904 | 71 | 71 | 0 | 17 (17-20) |
|  | FEX002 | NCT01340157 | 12 | 12 | 0 | 15 |
|  | FEX003 | NCT01483170 | 22 | 22 | 0 | 17 (11-17) |
| *g*-HAT | FEX004 | NCT01685827 | 203 | 125 | 78 | 6 (3-6) |
|  | FEX006 | NCT02184689 | 114 | 64 | 50 | 5 (4-5) |
| Chagas | CH-FEX001 | NCT02498782 | 40 | 12 | 28 | 3 (1-6) |

The formation of the M2 fexinidazole sulfone metabolite was modeled as a first-order “absorption” process. A one-compartment disposition model without inter-individual random variability and with additive error (on the log scale) provided the base model. Inclusion of inter-individual random variability in pharmacokinetic parameters provided a significant improvement. Inclusion of 1 and 2 transit compartments for the formation of the metabolite M2 also provided significant improvement to model fits as evidenced by the conditionally weighted residual plots. A two-compartment disposition model did not provide an improved fit to the data as quantified by a non-significant improvement in OFV. The final model was thus a one-compartment disposition model with two transit compartments for the formation of the metabolite. This structural model estimated that the metabolite M2 had a relative “absorption” (i.e. formation) rate of 0.04*h*^−1^/*F* (IIV of 26%), a volume of distribution of 80*L/F* (IIV of 20%) and a clearance of 3*L/h/F* (IIV of 14%).

### Liver toxicity in fexinidazole treated asymptomatic Chagas disease patients

In the dose ranging study of fexinidazole for the treatment of chronic asymptomatic Chagas disease, dose dependent elevations in AST and ALT were observed in a subset of patients. The time-series data of the liver transaminases elevations over the first 150 days following start of treatment show considerable heteroscedasticity but clear separation between patients with high and low M2 exposures (Fig 2). Some Chagas disease patients with exposures in the ranges seen in *g*-HAT (green lines) had liver transaminases elevated above 3 times the upper limit of normal (ULN). Peak elevations of ALT and AST were observed between 50 and 100 days after the start of treatment. For the patients whose levels rose above 3 times the ULN (for males: 53 and 46 units/L for ALT and AST, respectively; for females: 40 and 39 units/L for ALT and AST, respectively), the duration of elevated transaminases varied between 3 to 168 days for ALT and 8 to 40 days for AST. The duration of elevation was explained partially by total drug exposure (S2 Fig). All patients’ values eventually returned to normal.

**Figure 1:**
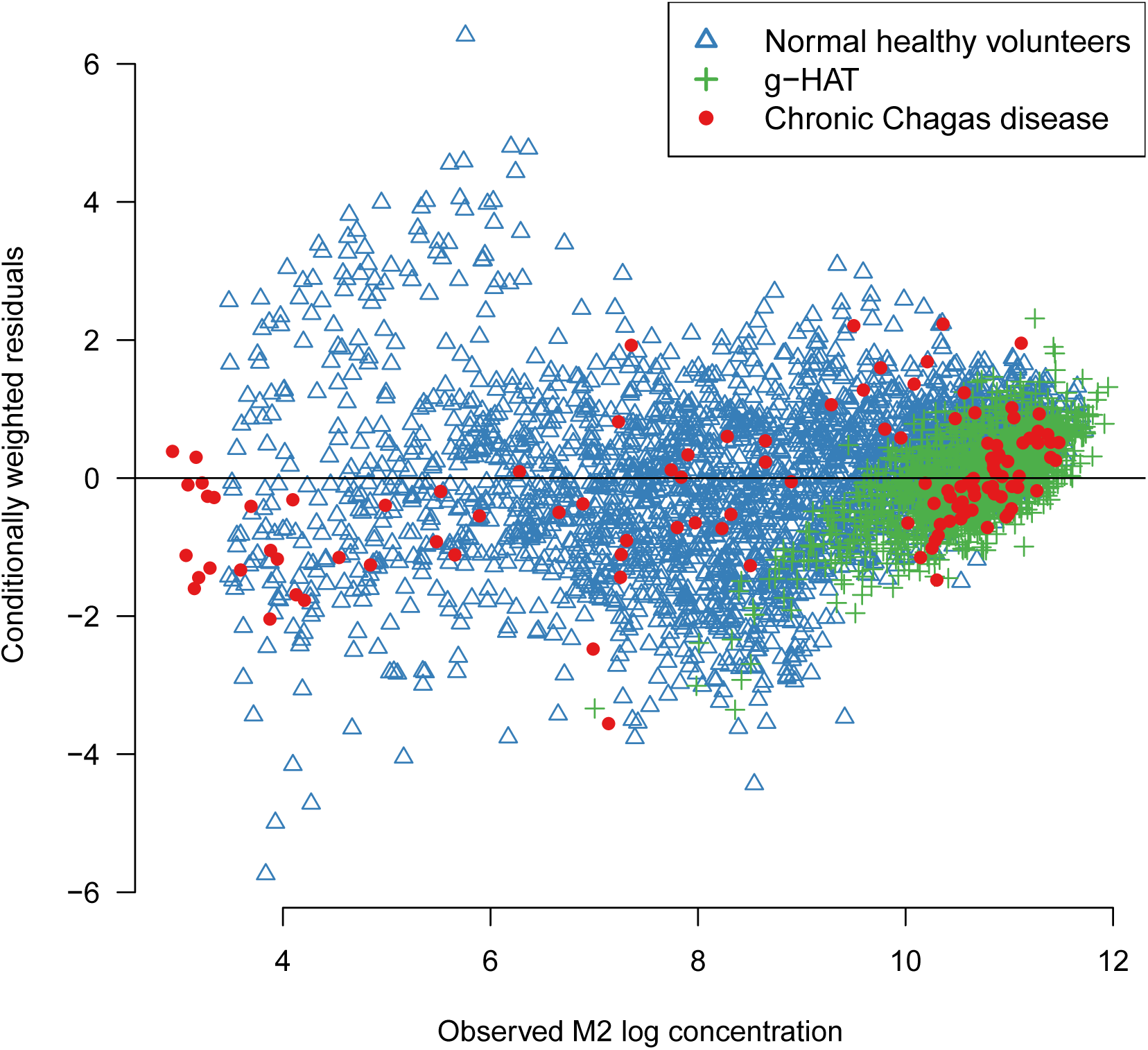
Visual pharmacokinetics model check. Comparison between the observed concentrations of the fexinidazole sulfone metabolite M2 and the conditionally weighted residuals of the final pharmacokinetic model fit to all M2 data from phase 1 trials (blue triangles), *g*-HAT sleeping sickness trials (green crosses) and asymptomatic Chagas disease trials (red circles).

**Figure 2:**
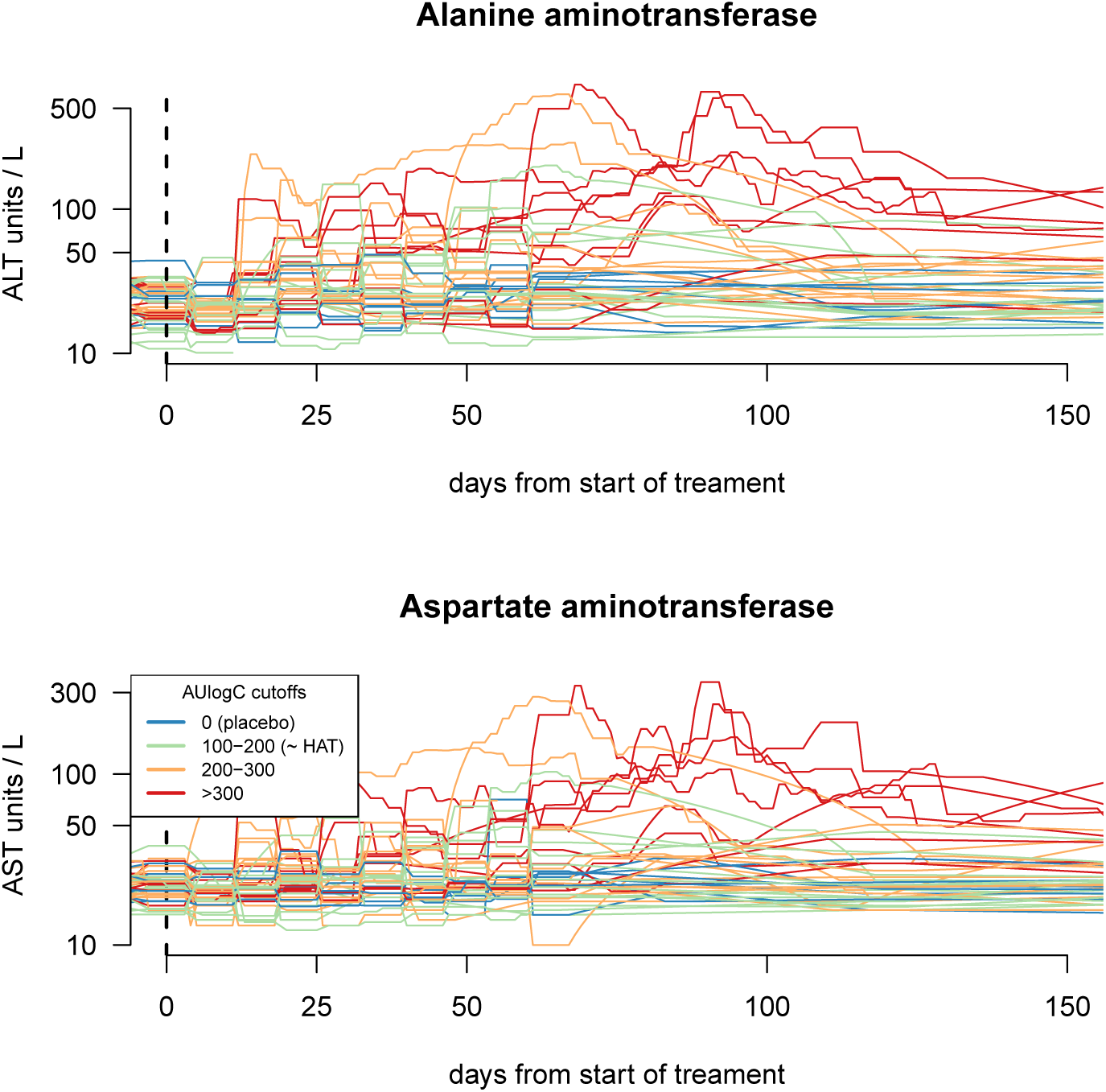
Time series data on the liver transaminases in all enrolled chronic Chagas patients. Top panel: ALT; bottom panel: AST. This shows data for all patients (n=47) enrolled in the trial of oral fexinidazole for the treatment of chronic Chagas disease. The absolute concentrations are shown on a log_10_ scale. In both panels, the colors correspond to individual total drug exposure as quantified by cumulative AUlogC of the M2 (sulfone) metabolite. Blue: placebo (no fexinidazole exposure); green: ranges of fexinidazole exposure corresponding to *g*-HAT regimen or less; orange: high ranges of AUlogC from 200-300; red: very high fexinidazole exposure with AUlogC above 300.

### Exposure dependent effect of fexinidazole on hepatic transaminases

For both ALT and AST, the posterior model fits indicate clear exposure-response relationships when the PD outcome is both measured as absolute peak increases (Fig 3, left column) and as relative fold increases (Fig 3, right column). As quantified by the *overlapping coefficient*, the posterior distributions give negligible probability (less than 5%) to the “null models” of no exposure dependent outcome in all four models (S4 Fig).

**Figure 3:**
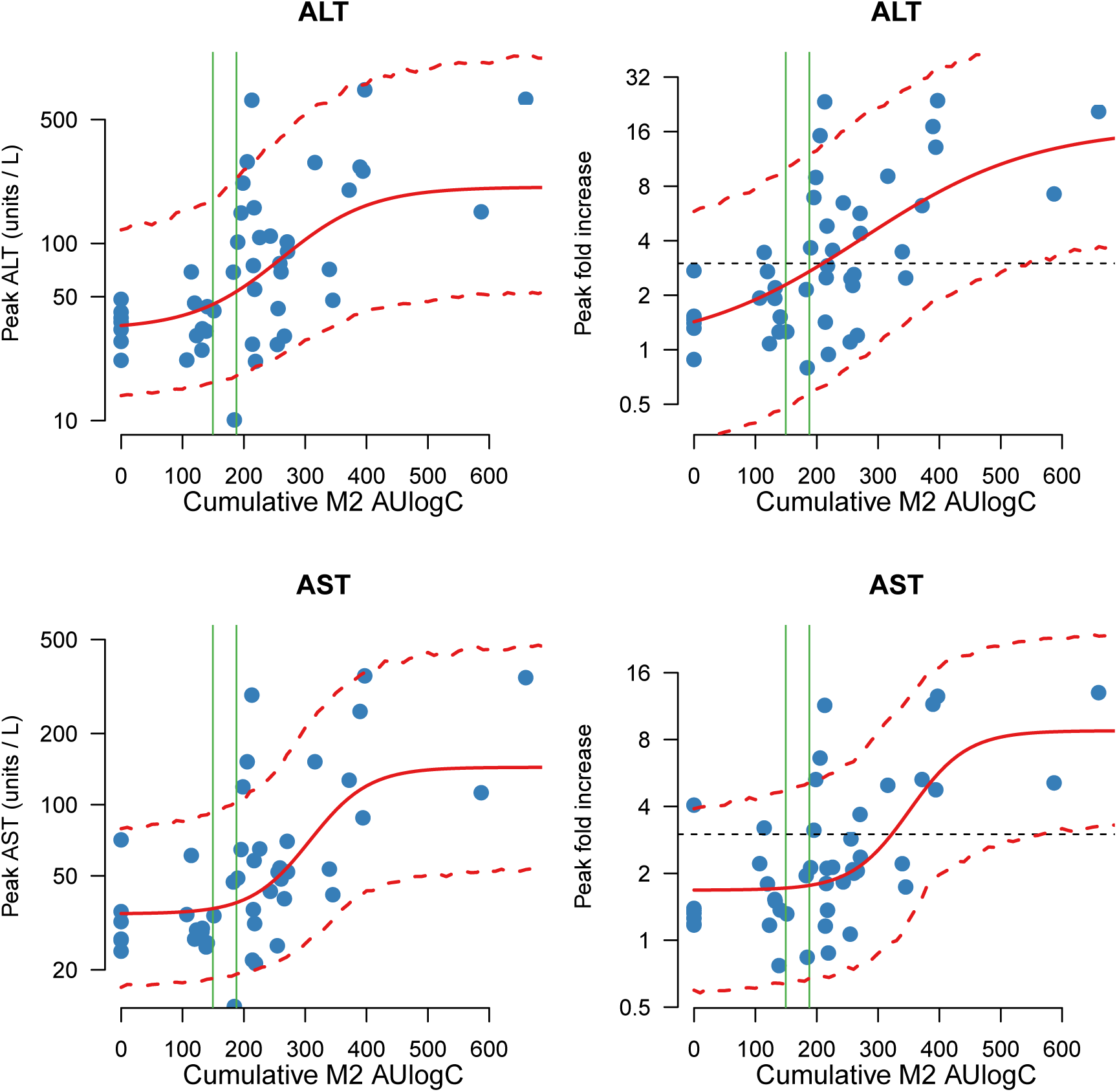
The exposure-response relationships for the peak liver transaminase values in fexinidazole treated chronic Chagas disease patients. Top row: ALT; bottom row: AST. The left column gives the relationship between fitted M2 AUlogC values and peak observed transaminase concentrations (y-axis is on the log_10_ scale). The right column gives the relationship between fitted sulfone metabolite (M2) AUlogC values and peak observed fold changes from baseline (y-axis is on the log_2_ scale). Individual data points are shown by the blue dots, sigmoid model mean fits along with 90% posterior prediction intervals are shown in red, and the range of AUlogC exposures with the *g*-HAT regimen is shown by vertical green lines. In the right column, the dashed black line shows the threshold value of 3 x baseline.

For ALT, the 80% credible intervals of the marginal posterior distribution over the *ED*_50_ overlap with the exposure intervals in the *g*-HAT regimen (Table 2). For AST, the 80% credible intervals of the marginal posterior distribution over the *ED*_50_ are above the exposure intervals in the *g*-HAT regimen (Table 2). The relationship between AUlogC and mg/kg dose in the Chagas trial is shown in Fig S1.

**Table 2:**
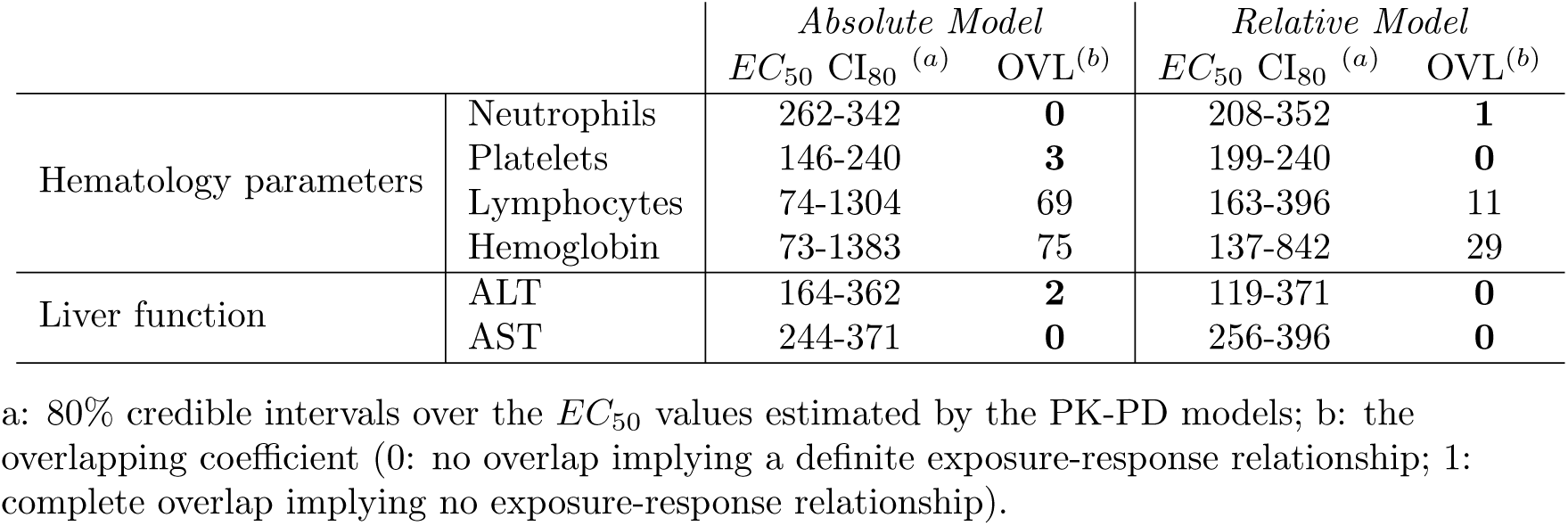
Summary of results from the Bayesian exposure-response models. Absolute models evaluate changes to the absolute values of the outcomes, and relative models evaluate fold changes from baseline in the values of the outcomes. Numbers in bold correspond to those with reasonable evidence of an exposure (dose) effect.

### Predicting liver toxicity in the *g*-HAT treatment regimen

The *relative* models fitted to data from asymptomatic Chagas disease predict that 50% of individuals with exposures (AUlogC) distributed as observed for the *g*-HAT regimen would have ALT elevations more than 2.8 times of baseline (more than a 180% increase), and AST elevations more than 2 times the baseline. However, no AST or ALT elevations greater than 2 times baseline were observed in any of the patients in the three field trials. Indeed, for ALT, the distribution of late measurements (all measurements taken between days 20-100) was identical to that of the early measurements (p=1, before and during treatment, Fig 4 bottom left panel). For AST, a significant difference was observed between early and late measurements (*p* < 0.01), but none had a fold change greater than 2 (Fig 4 bottom right). Therefore the model based on Chagas disease patients over-predicted liver toxicity substantially for both transaminases in *g*-HAT, indicating an effect specific to the Chagas disease population.

**Figure 4:**
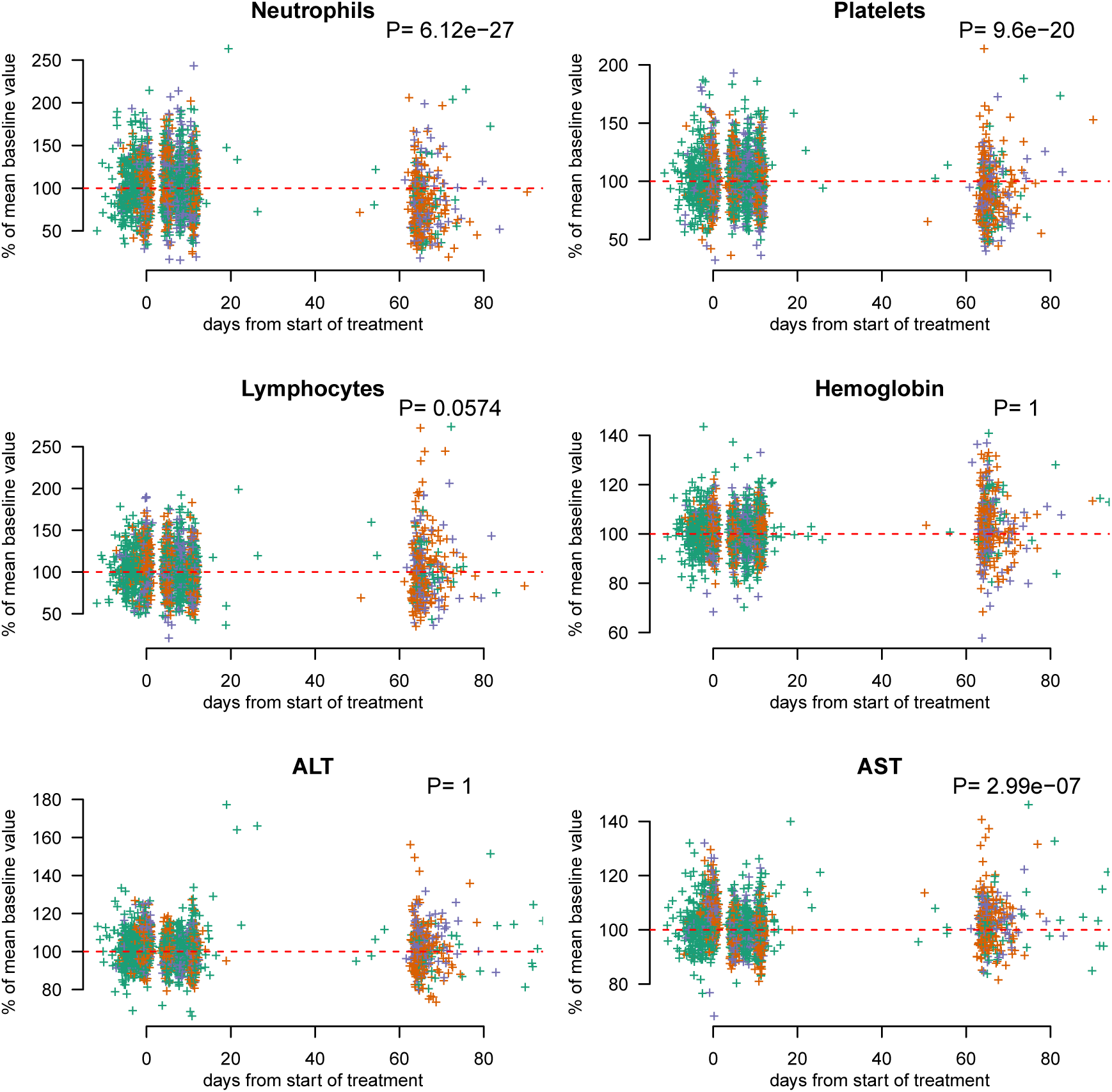
Pharmacodynamic outcomes in the *g*-HAT field trials. Each panel is a scatter plot of pharmacodynamic outcomes shown as % relative change from ‘baseline’ mean value as a function of time from start of treatment for all three field trials of fexinidazole for the treatment of *g*-HAT (*T. gambiense*). ‘Baseline’ is estimated as the average of all values taken up to day 20 post start of treatment. Colors correspond to trial ID. Green: randomized trial in adults with stage 2 *g*-HAT (FEX004), only fexinidazole treated individuals; orange: adults with stage 1 or 2 *g*-HAT (FEX005); purple: children with stage 1 or 2 *g*-HAT (FEX006). P-values are computed from a Mann-Whitney U test between the early (before day 20) and late (after day 20) groups of measurements. ALT: alanine aminotransferase concentration; AST: aspartate aminotransferase concentration.

## Hematology

### Hematological variables in fexinidazole treated chronic Chagas patients

In the dose-finding study of fexinidazole in chronic Chagas disease, 8 out of 40 patients who were assigned fexinidazole had reductions in neutrophil counts falling below 1000/*µ*L (compared to none in the placebo group). The day of the nadir value in this subgroup was day 65 (range: day 63 to day 71). These events were temporary with rapid recoveries. The median estimated duration of neutropenia (below 1000/*µ*L) calculated using linear interpolation between adjacent time points was 8.5 days (range: 7 to 21 days). All these patients had fexinidazole exposures as quantified by the AUlogC values above 300 (i.e. considerably greater exposures than seen in *g*-HAT, S1 Fig).

In addition to these more extreme variations, there was a consistent decrease in median neutrophil counts over the course of study. This trend began from the start of treatment up until the population nadir on day 70 in all patients not receiving placebo treatment (Fig 5, top left panel). The overall profile was one of a steady decrease with a greater reduction around two months after starting treatment. Day 57 was the median day of observed nadir in fexinidazole treated individuals and day 70 was the population median nadir value. These decreases were transient and a gradual recovery in neutrophil counts was observed in the follow-up visits.

**Figure 5:**
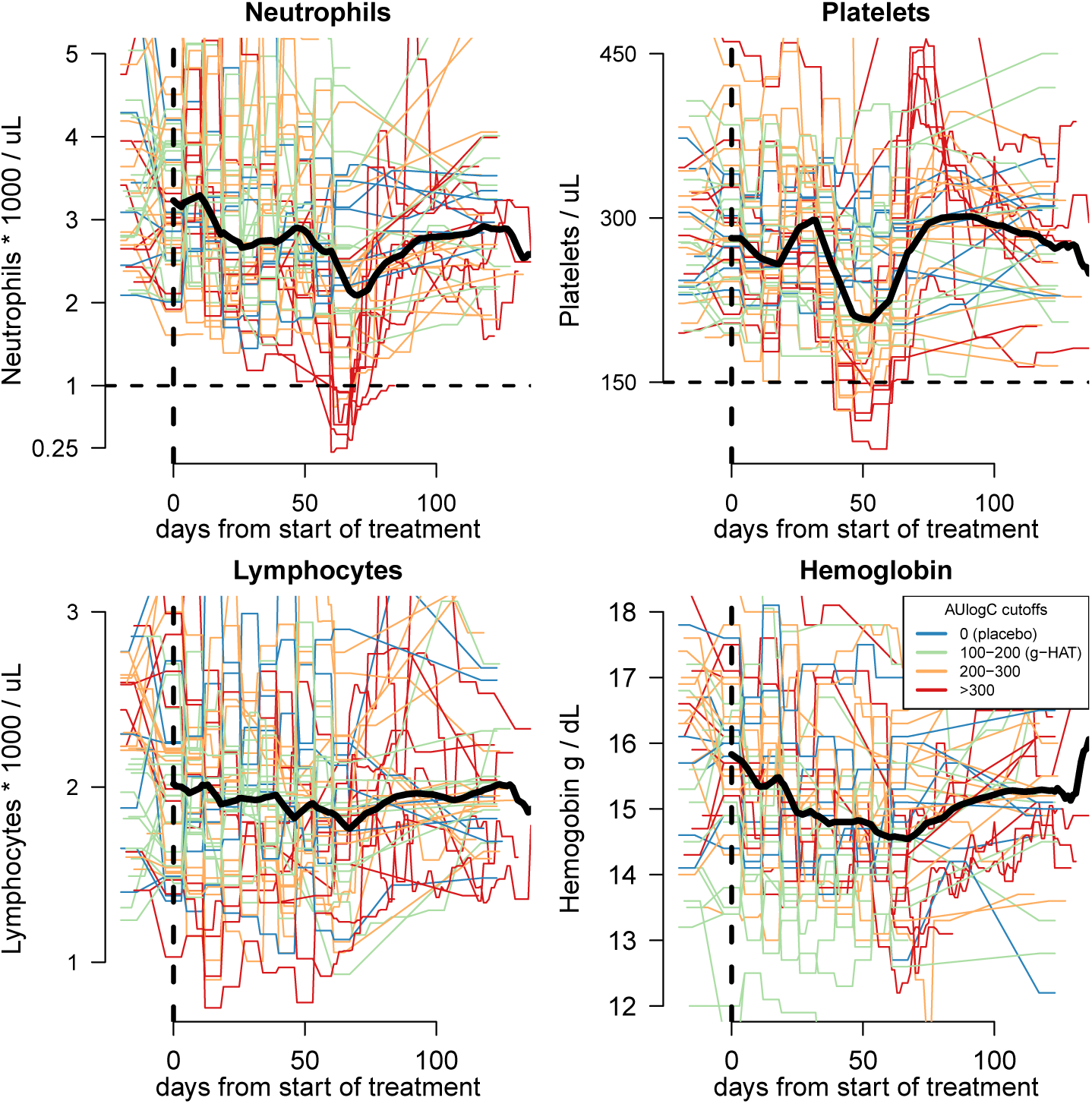
Time series data on hematological variables for all enrolled chronic Chagas disease patients. Start of treatment (day 0) is shown by the vertical black dashed line. Colors correspond to individual total fexinidazole exposure (AUlogC). The median trend for each hematological variable is shown by a thick black line. The threshold values of 1000 neutrophils per *µL* and 150 000 platelets per *µL* are shown by the horizontal black dashed lines.

The time series data for platelet counts show a similar temporal trend, albeit with a less marked initial reduction and a more marked later reduction (Fig 5, top right panel). All platelet counts remained above 50,000/*µ*L but 9 fexinidazole treated individuals had nadir counts below 150,000/*µ*L, 5 of whom were also in the neutropenia subgroup. The nadirs of neutrophil and platelet counts were correlated significantly; *ρ*= 0.5 (95%CI 0.2 to 0.7), p= 0.003. Nadir platelet counts were seen approximately two weeks before nadir neutrophil counts (median observed day of nadir in fexinidazole treated patients occurred on day 44; median population value was on day 53). In patients with platelets counts below 150,000/*µ*L, the median estimated duration of thrombocytopenia calculated using linear interpolation between adjacent time points was 9 days (range: 5 to 23 days).

Median hemoglobin counts had also dropped by 1.3 g/dL by day 67, the population median nadir value (Fig 5, bottom right panel). In contrast, lymphocyte counts showed no clear trend (Fig 5, bottom left panel).

### Hematological exposure-response effects of fexinidazole

For both neutrophil and platelet counts the models indicate clear exposure-response relationships when the PD outcome is measured both as absolute peak increases (Fig 6, left column) or as relative fold increases (Fig 6, right column). As quantified by the *overlapping coefficient*, the posterior distributions give negligible probability (less than 5%) to the “null model” of no exposure dependent outcome in all four models (S4 Fig. For lymphocyte counts and hemoglobin concentrations, there is not clear evidence of an exposure-response effect, with OVL coefficients varying between 11 and 75%.

**Figure 6:**
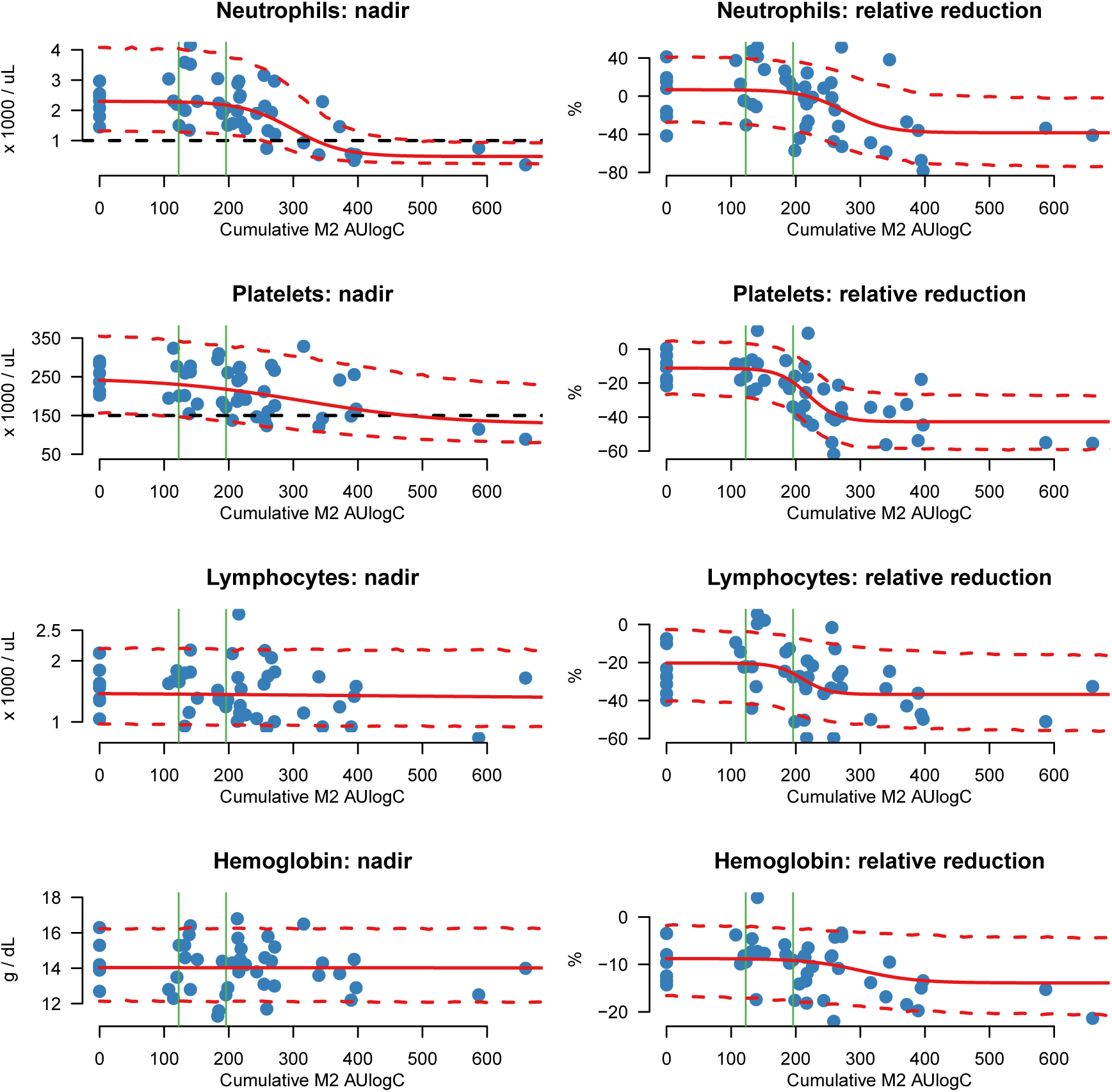
The exposure-response relationships for the main hematological variables of interest in the dose finding assessment of fexinidazole in asymptomatic Chagas disease. Exposure is quantified by the total cumulative M2 metabolite AUlogC. The left column shows the scatter plots between fitted exposures and nadir observed values (blue dots). The right column shows the scatter plots between M2 AUlogC exposure and the maximum observed relative % decrease with respect to the individual baseline value. Sigmoid model mean fits along with 90% posterior prediction intervals are shown in red. The range of fitted drug exposures after the *g*-HAT regimen are shown by the vertical green lines.

For reductions in neutrophil counts, the credible intervals over the *EC*_50_ parameters in both the *absolute* and *relative* models are above *g*-HAT exposures (Table 2). However for the reductions in platelet counts, the absolute model suggests an *EC*_50_ within the *g*-HAT exposures (Table 2).

### Predicting drug effects on hematological variables in *g*-HAT

For exposures distributed according to those observed in the *g*-HAT regimen, the *relative* models trained on the data from the chronic Chagas patients predicted a median relative decrease in neutrophil counts of 0%, with 10% of patients predicted to have relative reductions of approximately 35%. For platelet counts, the predictions were for a median decrease of 20% with 10% experiencing decreases of more than 35%.

Statistically significant but clinically insignificant decreases from baseline were observed in both neutrophil and platelet counts in *g-*HAT patients (*p* < 0.001 for both; Figure 4: top two panels). The median decreases were 20% and 10% for neutrophils and platelets, respectively. The nintieth percentiles were 60% & 40%, respectively. The timing of the late full blood counts done in the *g*-HAT trials coincided with the timing of the population nadir for neutrophil counts in the Chagas trial but not the population nadir for platelet counts (two weeks earlier). This difference in the timing of hematological changes was not known at the time of the *g*-HAT trial protocol amendments.

These observed reductions in neutrophil and platelet counts could be confounded. Patients in the Chagas study were asymptomatic, whereas *g*-HAT patients were ill and thus may have had higher neutrophil and platelet counts on admission. This possible confounding effect can be approximated by comparing the observed decreases in the *g*-HAT patients receiving fexinidazole and those receiving NECT in the FEX004 study (randomized assignment). For both platelet and neutrophil counts, there were no significant reductions for the NECT group whereas there were in the fexinidazole group (Figure 7).

**Figure 7:**
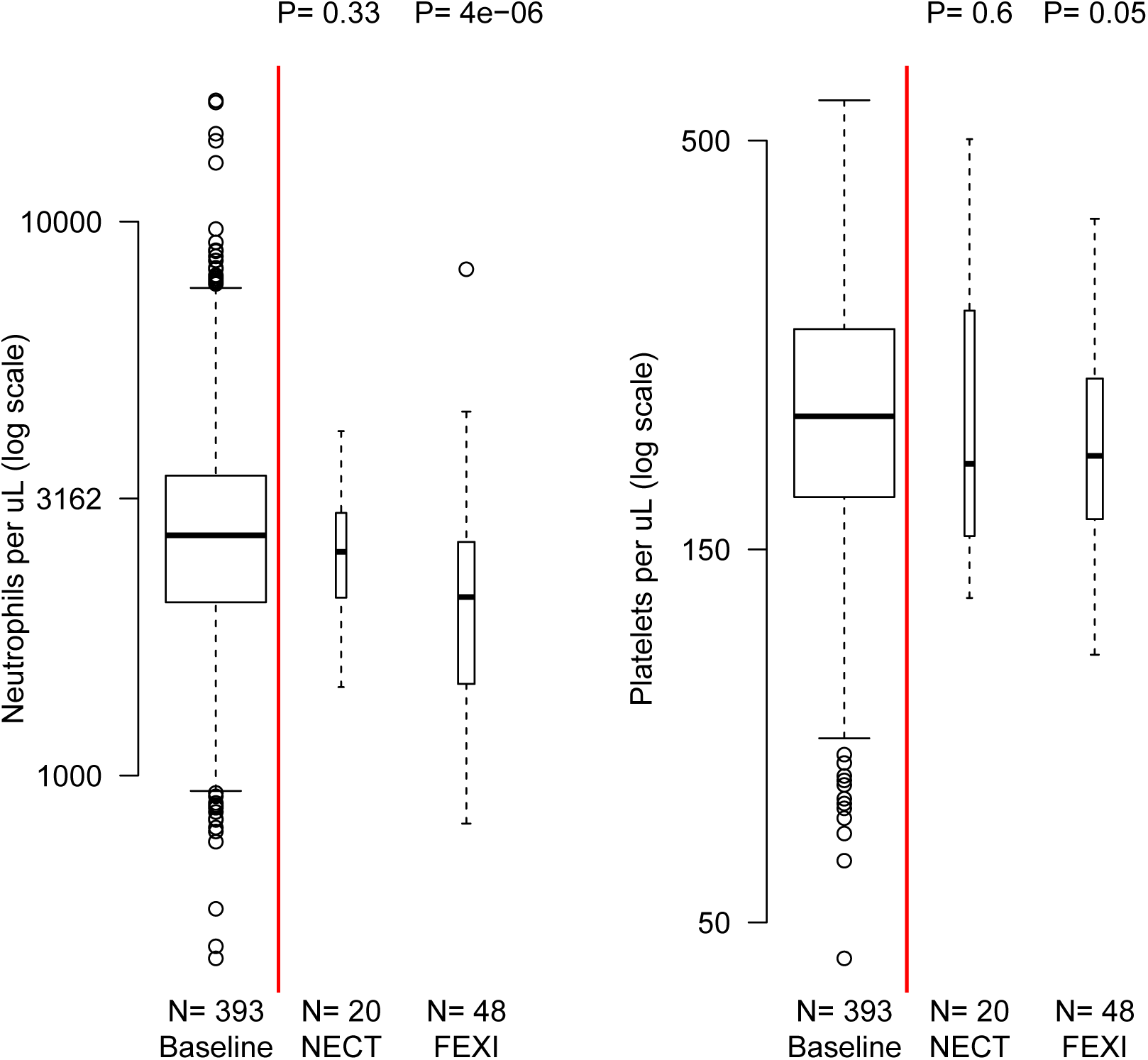
Comparison between early (baseline and during treatment) and late (circa day 70) neutrophil (left panel) and platelet counts (right panel), grouped by randomised treatment in trial FEX004 (late stage adults with *g*-HAT). P-values are computed from a Mann-Whitney U test between early (all patients) and late measurements. The width of each box-and-whisker plot is proportional to the square root of the number of observations.

Although the numbers were small (*N* = 20 late counts in the NECT arm), this suggests that the fexinidazole regimen for *g*-HAT treatment results in mild but predictable delayed reversible decreases in neutrophil and platelet counts.

## Discussion

Fexinidazole has the potential to replace nifurtimox-eflornithine as the treatment of choice for human African trypanosomiasis (*g*-HAT) (1; 2; 3; 15). In large, well controlled studies, it has proved well tolerated and effective (findings from FEX004 are reported in (9) and findings for FEX005 & FEX006 are as yet unpublished). A safe, once daily, oral treatment would substantially improve the prospects for elimination of this major tropical neglected disease. These excellent clinical results in *g*-HAT and the significant *in vitro* activity of fexinidazole against other kinetoplastid parasites prompted investigations in leishmaniasis and Chagas disease, but the preliminary investigations in chronic indeterminate Chagas disease were interrupted when some patients developed severe but transient delayed neutropenia. In addition significant elevations in aspartate and alanine aminotransferases were noted suggesting liver toxicity. These adverse reactions were unusual in that they occurred sometimes up to two months after starting the fexinidazole treatment, often well after the drug had stopped and the bioactive parent and metabolites would have been cleared.

Fexinidazole is converted *in vivo* to sulfoxide and sulfone metabolites which retain biological activity, but the majority of bioactive exposure *in vivo* is to the sulfone (M2) metabolite. It was not possible to dissociate toxicity relationships of the parent compound and metabolites and so the only associations explored here are with the sulfone metabolite. The mechanisms of fexinidazole toxicity are not known but appear to be class effects, although why neutrophil and platelet counts fall approximately nine and seven weeks after starting, respectively, and in many cases weeks after completing fexinidazole treatment, is not known. The sequential timing of neutrophil and platelet reductions and their correlation without coincidental lymphopenia suggests bone marrow suppression. The transient nature of these reductions (lasting approximately one week) suggests a transient inhibition or suppression of a bone marrow precursor.

This pharmacokinetic-pharmacodynamic assessment used total cumulative log concentrations (total AUlogC) of the metabolite M2 as a proxy measurement of drug exposure. This is justified for several reasons. First, the ease of estimation as M2 is formed slowly and has the most predictable pharmacokinetic behavior with the least inter-individual variability. Thus in the dose assessment trial of fexinidazole in Chagas disease which provides the large majority of the pharmacodynamic information, sparse drug measurements taken over 8 weeks can be summarized reliably by model estimates of M2 AUlogC. Secondly, M2 concentrations have been shown to be the best predictor of electrocardiograph QT prolongation (unpublished observations) and vomiting. The relationship between the total mg/kg dose of fexinidazole and the pharmacokinetic exposure is imprecise for higher doses but gives an approximate threshold dose of 400 mg/kg for a threshold AUlogC exposure of 300 (S1 Fig).

The nitroimidazoles are a class of drugs with known potential for both neutropenia and liver toxicity (16; 17; 18; 19; 20; 21), although cholestatic hepatitis is the usual manifestation (22). One patient in the phase 1 studies of fexinidazole showed evidence of hepatotoxicity on day 15 after 14 daily doses of 3600mg. Values returned spontaneously to normal (6). No case meeting Hy’s Law criteria was reported. However, follow-up in the phase 1 studies was short (a maximum of 28 days post start of regimen) so later asymptomatic hepatotoxicity following these studies cannot be excluded. The duration of the dose regimens of fexinidazole evaluated in Chagas disease were based on the regimen of benznidazole, another nitroimidazole which is the currently recommended treatment of Chagas disease and which also has been associated with both hepatotoxicity and neutropenia (23). Benznidazole is usually given for eight weeks, and so in some patients total mg/kg dosing of fexinidazole was substantially higher than in the 10 day *g*-HAT regimen (in one arm it was 7 times higher). The principal findings of this retrospective PK-PD study are that the risks of both increased transaminases, delayed neutropenia, and delayed platelet reductions are proportional to drug exposure, but that exposures in *g*-HAT are below those associated with clinically significant toxicity. The estimated EC50 quantified in terms of total M2 AUlogC for hepatoxicity, and reductions in neutrophil and platelet counts are considerably higher than the maximum observed M2 AUlogC values in patients treated for *g*-HAT.

The observed reductions in neutrophil and platelet counts in the *g*-HAT field trials of fexinidazole match predictions from the PK-PD model built using the data from the dose assessment trial in Chagas disease. These were mild and clinically insignificant reductions in neutrophil and platelet counts which are very unlikely to pose a threat to *g*-HAT patients. On the other hand, the predicted dose-dependent hepatotoxicity based on the Chagas disease data was not observed in the treatment of *g*-HAT by fexinidazole. This suggests an additional disease effect which is specific to Chagas disease, or to the Bolivian population studied. Future studies of fexinidazole for the treatment of Chagas disease need to have long follow-up (more than six months) in order to assess potential iatrogenic changes to hematological variables and liver transaminases.

## Conclusion

Taken together these data suggest that there is a satisfactory margin of safety for dose related toxicity with the current fexinidazole regimen for the treatment of *g*-HAT. Future trials of fexinidazole in Chagas disease should assess liver function over a period of at least six months following the start of treatment. Shorter regimens of fexinidazole (less than ten days) should be safe for the treatment of chronic Chagas disease. Transient but clinically significant neutrophil decreases are to be expected in individuals taking a total dose of more than 400mg/kg of oral fexinidazole.

## Materials and methods

### Clinical trials

#### Phase 1 studies in normal healthy volunteers

Three separate phase 1 studies were carried out in 116 normal healthy male volunteers (NHV) to assess the tolerability of oral fexinidazole and to characterize its pharmacokinetics. Full details of these phase 1 studies and results from non-compartmental pharmacokinetic analyses are reported elsewhere (6). Electrocardiographic recordings were collected in all trials, but these data are being analyzed separately.

##### FEX001

This was a randomized, double-blind, placebo controlled study of the tolerability and the pharmacokinetics of oral fexinidazole given in single and repeated doses (clinical trial ID: NCT00982904). This study also included a comparative bioavailability study of an oral suspension versus the tablet formulation, and an exploratory assessment of food effects. All subjects were healthy male volunteers of sub-Saharan origin. The pharmacokinetics of oral fexinidazole were characterized in three separate sub-studies. Dense pharmacokinetic sampling was performed in all sub-studies at the following time points: pre-dose, 0.5, 1, 2, 3, 4, 6, 9, 12, 16, 24, 48, 72, 96, 120, 144 & 168 hours post-dose.

- Part 1: single doses ranging from 100 to 3600mg (n=54).
- Part 2: testing for bioequivalence between the oral tablet form and the oral suspension form which was not included in this analysis (n=11).
- Part 3: daily dosing for 14 consecutive days with doses of 1200, 2400 & 3600mg (n=17).

##### FEX002

This was a randomized, open label study to assess the effect of two different types of food versus fasted conditions on the relative bioavailability of a single dose of oral fexinidazole in healthy males (n=12) (clinical trial ID: NCT01340157). Free fraction plasma concentration measurements were taken at 5 nominal time-points (1, 4, 12, 24, & 72 hours post-dose). Frequent venous sampling was done at the same time-points as for the FEX001 study.

##### FEX003

This was a randomized, double blind, placebo controlled comparison of two 10 day regimens of fexinidazole (clinical trial ID: NCT0148370) both including a 4 day loading dose. Regimen 1 was administered as 1800mg for 4 days, followed by 1200mg for 6 days. Regimen 2 was administered as 2400mg for 4 days followed by 1200mg for 6 days. Each dose was taken after a meal. All subjects (*n* = 22 completed the study out of 30 included subjects) were male healthy volunteers with both parents of sub-Saharan African origin. Pharmacokinetic samples were taken on days 1, 4 & 7 at the following nominal time-points: pre-dose and then 0.5, 1, 2, 3, 4, 6, 9, 12, 16 & 24 hours post-dose. On day 10 (last dose), pharmacokinetic samples were taken pre-dose and then 0.5, 1, 2, 3, 4, 6, 9, 12, 16, 24, 48, 72, 96, 120, 144 & 168 hours post-dose.

#### Treatment trials in *T.b. gambiense* sleeping sickness (*g*-HAT)

Phase 2/3 trials were conducted in the Democratic Republic of Congo (DRC) and Central African Republic (CAR). The pivotal study was conducted in 394 adult stage 2 *g*-HAT patients (i.e. in patients with CNS involvement). Two additional cohort studies were conducted in 230 patients with stage 1 *g*-HAT (i.e. no CNS involvement) and 125 children with both stages of *g*-HAT who were aged 6-14 and weighed more than 20kg. All patients or their guardians provided full informed consent. All patients had parasitologically confirmed *g*-HAT and a Karnovsky score > 50 in order to be eligible for enrollment. Adult patients and children weighing more than 35kg were treated with fexinidazole 1800mg once daily for 4 days followed by 1200 mg once daily for 6 days. Children older than 6 years and weighing between 20 and 35kg were given an adapted regimen: 1200mg for 4 days followed by 600mg for 6 days. All field trials administered fexinidazole as 600mg tablets in blister packaging.

##### FEX004

This was the pivotal Phase 2/3 study assessing the safety and efficacy of fexinidazole in the treatment of *g*-HAT (clinical trial ID: NCT01685827) (9). It was an open label randomized trial of oral fexinidazole compared to the current standard-of-care regimen of Nifurtimox-Eflornithine Combination Therapy (NECT) in adult patients (> 15 years old) with late-stage *g*-HAT. Late stage sleeping sickness is defined as confirmed parasites in the cerebrospinal fluid (CSF) or confirmed parasites in blood and a CSF white cell count greater than 20/*µ*L. The randomization ratio was 2:1 with 264 patients enrolled into the fexinidazole arm and 130 patients enrolled into the NECT arm. The NECT regimen was a combination of oral nifurtimox tablets, 5 mg/kg three times daily for 10 days (D1 to D10); and eflornithine 200 mg/kg administered twice daily as a 2-hour IV infusion for 7 days. The fexinidazole adult regimen was 4 daily doses of 1800mg followed by 6 daily doses of 1200mg. This is referred to as the *g*-HAT regimen. The study took place at 10 sites in the Democratic Republic of Congo and the Central African Republic.

Pharmacokinetic samples were taken in 203 fexinidazole treated patients. The field sites were very remote and it was not possible to store frozen samples on-site. Capillary whole blood was therefore collected and aliquoted onto filter paper to produce dry blood spots (DBS: 300*µ*L (10)) from a finger-prick sample at the following time points: day 8: 3 hours after dose; day 9: 3 hours after dose; day 10: 3 hours and 7 hours after dose; days 11 and 12: 24 and 48 hours after the last dose. A lumbar puncture was performed on day 11 (24 hours after the final dose) for the efficacy assessment. In some patients an aliquoted (300*µ*L) CSF sample from the follow up lumbar puncture was allowed to dry on filter paper and stored for later drug measurement (n=82). Full blood counts and biochemistry were taken at enrollment and then on days 5 and 11 after the start of treatment. In response to the hematology and biochemistry abnormalities recorded during the asymptomatic Chagas disease treatment trial, the protocol was modified to allow for a small subgroup of patients (*n* = 68) to have full blood counts and biochemistry checked 9 weeks after the start of treatment.

##### FEX005

This was a first “plug-in” study which included stage 1 and early stage 2 adult *g*-HAT patients receiving the same fexinidazole regimen as FEX004 (clinical trial ID: NCT02169557). It was an open label single-group study which enrolled 230 patients. No pharmacokinetic sampling was carried out, but full blood counts and biochemistry were taken at enrollment and then on days 5 and 11 and at 9 weeks following the start of treatment.

##### FEX006

This was a second plug-in study of fexinidazole in children older than 6 years old and weighing more than 20kg with stage 1 and 2 *g*-HAT (clinical trial ID: NCT02184689). Children weighing between 20 and 35kg were given 1200mg of fexinidazole daily on days 1-4 (2/3 of the adult dose) and then 600mg daily on days 5-10 (half the adult dose). Children weighing more than 35kg were given the adult regimen. This analysis included pharmacokinetic data from 114 patients. A series of pharmacokinetic measurements using DBS were taken on days 10 (3 and 7.25 hours after the final dose), 11 (24 hours post final dose) and 12 (48 hours post final dose). Drug measurement was also performed on dried CSF on filter paper (as in FEX004) from the day 11 lumbar puncture in the first 30 patients.

#### Treatment trial in chronic Chagas disease

A dose finding study (clinical trial ID: NCT02498782) in adult Bolivian patients with chronic indeterminate Chagas disease (referred to as chronic Chagas) began in July 2014. Non-pregnant adult patients were enrolled if they were positive for a validated *T. cruzi* PCR test, but had no clinical evidence of end organ damage. The duration of treatment was structured around currently recommended regimens for the treatment of *T. cruzi* infections with benznidazole. All patients were outpatients. Patients were advised to take the treatment as a single daily dose and with a meal. Each week they were given enough medication until the next scheduled weekly visit. Treatment was thus unobserved and drug adherence was checked weekly by pill counting. Benznidazole rescue treatment at the end of study was offered for non-responders.

Patients were randomized to one of seven once-daily dosing regimens:

1. Fexinidazole 1200mg daily for 2 weeks followed by matching placebos for 6 weeks.
2. Fexinidazole 1200mg daily for 4 weeks followed by matching placebos for 4 weeks.
3. Fexinidazole 1200mg daily for 8 weeks.
4. Fexinidazole 1800mg daily for 2 weeks followed by matching placebos for 6 weeks.
5. Fexinidazole 1800mg daily for 4 weeks followed by matching placebos for 4 weeks.
6. Fexinidazole 1800mg daily for 8 weeks.
7. Placebos for 8 weeks.

After completion, patients were followed for 12 months. Pharmacokinetic samples were taken on day 0 (pre-dosing), on day 1, and then weekly for weeks 2-5 and then on weeks 9 & 10. Laboratory hematology and biochemistry samples were taken on day 0, weekly for weeks 2-10. The study was interrupted approximately three months later after the enrollment of 47 subjects due to toxicity concerns.

### Drug measurements

Blood samples from the phase 1 studies were taken into lithium heparin tubes, centrifuged immediately and plasma was separated and stored at −70^*◦*^C until bioanalysis. Fexinidazole and its sulfoxide (M1) and sulfone (M2) metabolites were analyzed on a Supelco Ascentis Express C18, 2.7*µ*m, 50 *×* 4.6mm I.D. column using a validated liquid chromatography–tandem mass spectrometry (LC-MS/MS) method. The plasma lower limit of quantification for fexinidazole, M1 and M2 was 0.5, 10 and 10ng/mL, respectively. For clinical sampling, Whatman #903 Protsaver 5 spot paper purchased from GE Healthcare Bio-Sciences (France) was used. Upon arrival of the DBS samples in the bioanalytical laboratory and before analysis, a visual inspection of the quality of the spot was performed. A blood spot was considered valid if the following criteria were met: (i) spot diameter was equal or greater than 7mm; (ii) spot was spread symmetrically on both sides of the sampling paper; and (iii) spot was made from a single drop of blood and was dark red in color. The blood volume deposited onto the filter paper and position of the punch had no effect on the quantitation of the test compound. Hematocrit values between 30% and 50% were shown not to affect the accuracy of drug measurement.

The ratio of capillary blood fexinidazole, M1 and M2 metabolites to simultaneous plasma concentrations stayed constant over time at 0.59, 1.00 and 0.97, respectively, regardless of the plasma concentration of fexinidazole, M1, M2 and sampling time.

### Pharmacokinetic analysis

The majority of the pharmacodynamic events of interest (evidence of hematological and liver related toxicities) occurred in the dose finding assessment of fexinidazole in chronic Chagas disease patients. This trial had only sparse pharmacokinetic sampling. Thus the primary goal of the pharmacokinetic modeling exercise was to impute as reliably as possible the pharmacokinetic profiles of these patients. Previous work (in press) has shown that the metabolite M2 is the primary determinant of both efficacy and adverse events (QT interval prolongation). For this reason we used the pharmacokinetic profile of the slowly eliminated sulfone metabolite (M2) as the determinant of the exposure related adverse events. This metabolite is also the most stable amongst the three compounds, exhibiting the least inter-individual variability (6).

The pharmacokinetic data analysis was done using NONMEM V.7.4 (ICON Development Solutions, Ellicott City). Molar units of the metabolite M2 concentrations were transformed into their natural logarithms and modeled using both 1 and 2 compartment disposition models with first order formation and elimination. Multiple candidate structural models were evaluated for the formation of the drug, using 0, 1, 2, or 3 transit compartments. The different structural models were evaluated on the dense data from the phase 1 studies. Scaling of parameters by weight was evaluated using the allometric relationship: 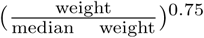 for clearances and a linear relationship (ratio of weight to median weight) for volumes. A food-effect was also introduced as a covariate for volume of distribution (in some phase 1 studies the drug was given to fasting subjects whereas in all field trials the drugs were administered after food). The final estimation of the model parameters used all data from Phase 1, *g*-HAT and Chagas field trials. Variation between these phase 1 and treatment trials (study and disease effects) were considered only as changes to the relative “absorption” parameter F (scaling parameter on F). The data from both the *g*-HAT and Chagas field trials are too sparse to estimate changes in absorption rate, clearance, or volume accurately. Trial specific effects were introduced in the model as categorical covariates with a linear effect in terms of percentage reduction for both clearance and volume parameters. Three categories were defined (NHVs, *g*-HAT trials, and Chagas trial). The final NONMEM model code is provided in the supplementary materials (S5 Code).

### Pharmacodynamic and statistical analyses

All the pharmacodynamic and statistical analyses were performed using R software (R Core Team 2016).

#### AUlogC to quantify drug exposure

From the dose finding study in Chagas disease the main pharmacodynamic events identified were delayed reductions in neutrophil counts and rises in plasma concentrations of liver transaminases. In pre-clinical and clinical studies there was no evidence for very slowly eliminated metabolites. Therefore, because of the long interval between drug exposure and these pharmacodynamic outcomes (i.e. after almost complete elimination of the drug and its active metabolites), the pharmacokinetic driver cannot be the concentrations at the time of the adverse effects but an overall summary of exposure with hysteresis in the concentration-effect relationship. In this work we use the total cumulative area under the log concentration curve (AUlogC) as the pharmacokinetic proxy for drug exposure.

#### Estimating exposure-response curves

This was a retrospective exploration of the relationships between total drug exposure and the main observed adverse events. All PK-PD models were fitted in a Bayesian framework using *stan* (11) with weakly informative priors for all parameters. Posterior distributions are given in the supplementary materials S3 Fig & S4 Fig. Model code and exact prior specification are provided in the supplementary materials (S6 Code).

Six pharmacodynamic outcomes were examined, four hematological (blood neutrophil counts, platelet counts, lymphocyte counts, and hemoglobin), and two related to liver toxicity (serum AST and ALT fold changes). The “steady state” dynamics of these four hematological parameters are substantially different. In the absence of biological perturbations, the red cell count and thus the blood concentration of hemoglobin is a very stable process within an individual, with daily variations of around *±.*5 g/dL. For individuals in the Chagas study, this corresponds to variations of approximately 3% around baseline. Neutrophil, platelet and lymphocyte counts exhibit much larger variations at steady state, with baseline intra-individual variation of up to 50% in this dataset. For this reason, quantifying time dependent changes to these four processes necessitates different methodologies. For example, large variations of *±*50% in neutrophil counts above a lower threshold count of 1000*/µL* are considered normal, whereas a decrease of 10% in hemoglobin could be considered clinically relevant even when staying within the “normal range” bounds. In order to account for these differences in temporal dynamics, we analyzed both the absolute values (nadir values for hematology and peak values for liver transaminases observed after start of treatment) and the relative values (maximum relative decreases or increases from baseline, respectively). Individual baseline values were calculated as the mean value observed in the pre-treatment visit and the day 0 visit (start of fexinidazole treatment), i.e. two values per person. For transparency we present results from both analyses for all pharmacodynamic outcomes. Throughout, models based on the absolute PD values are denoted the *absolute models*, and those based on the relative changes are denoted the *relative models*.

To estimate the exposure-response curves for the hematological parameters and the liver toxicity outcomes, we fitted four parameter sigmoid functions, defined as:

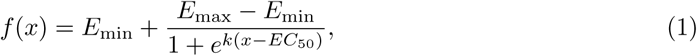

where *f* is the pharmacodynamic outcome of interest modeled as a function of drug exposure *x*; *E*_min_ is the baseline mean outcome under no or negligible drug exposures; *E*_max_ is the asymptotic maximal effect for high drug exposures; *EC*_50_ is the drug exposure corresponding to 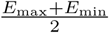 (half-maximal effect concentration); and *k* parameterizes the slope of this sigmoid relationship. In the *stan* specification of the model, the slope parameter *k* was explored in log space to improve convergence.

The *relative models* (which can have both positive and negative outcome values) use an normal additive error term parameterized by its standard deviation *σ_add_*. The *absolute models* (only positive outcomes) use a proportional normal error term *σ_prop_*: *log*(*y*) *∼ N* (*f* (*x*)*, σ_prop_*), where *y* is the observed outcome and *f* (*x*) is the sigmoid regression mean prediction.

#### Post-hoc evaluation of an exposure-response relationship

This was a post-hoc PK-PD analysis and as such was prone to data dependent analyses and false positive results (12). We attempted to minimize this danger by avoiding an analysis contingent on significant p-values of a null hypothesis that there is no exposure-response relationship (which can be difficult to control properly for multiple comparisons) and instead summarized the posterior evidence for exposure dependent PD outcomes.

The “null model” for equation 1 is one for which there is no exposure dependent PD outcome, and can be defined as the model for which *E*_min_ = *E*_max_. Thus the model collapses to a simple zero-order linear trend (slope coefficient of zero). The posterior evidence of the “null model” is quantified by the overlap between the marginal posterior distributions over *E*_max_ and *E*_min_. This overlap is quantified by the *overlapping coefficient*, defined as a distance metric between two densities *f* and *g* with the same support (13):

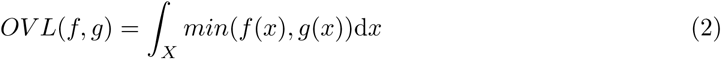

This is an intuitive measure of the overlap between two arbitrary densities of same support *f* and *g*, with values varying between 1 (complete agreement) and 0 (complete disagreement).

#### Predicting toxicity in the fexinidazole regimen recommended for *g*-HAT

The trial of fexinidazole in chronic Chagas disease had regular full blood counts (weekly up to week 10, then monthly) providing reasonable confidence that the observed time-series patterns for hematological parameters and liver function are indicative of the true underlying patterns. For the three *g*-HAT field trials, weekly blood counts in the weeks following treatment were not performed. Some late measurements were taken in FEX004 (late stage adults), and most patients in the FEX005 and FEX006 trials had one late measurement (around week 10). Thus the distribution of nadir relative decreases for neutrophil counts is expected to be different between the two studies because of the trial design. Because of the biases introduced by these different trial designs and the differences in the two populations (for example, individuals of sub-Saharan African descent have lower neutrophil counts due to different genetic variants of the Duffy gene (14)) we cross predicted using only the *relative* models (nadir or peak values divided by baseline value).

## Supporting information

**S1 Fig.**
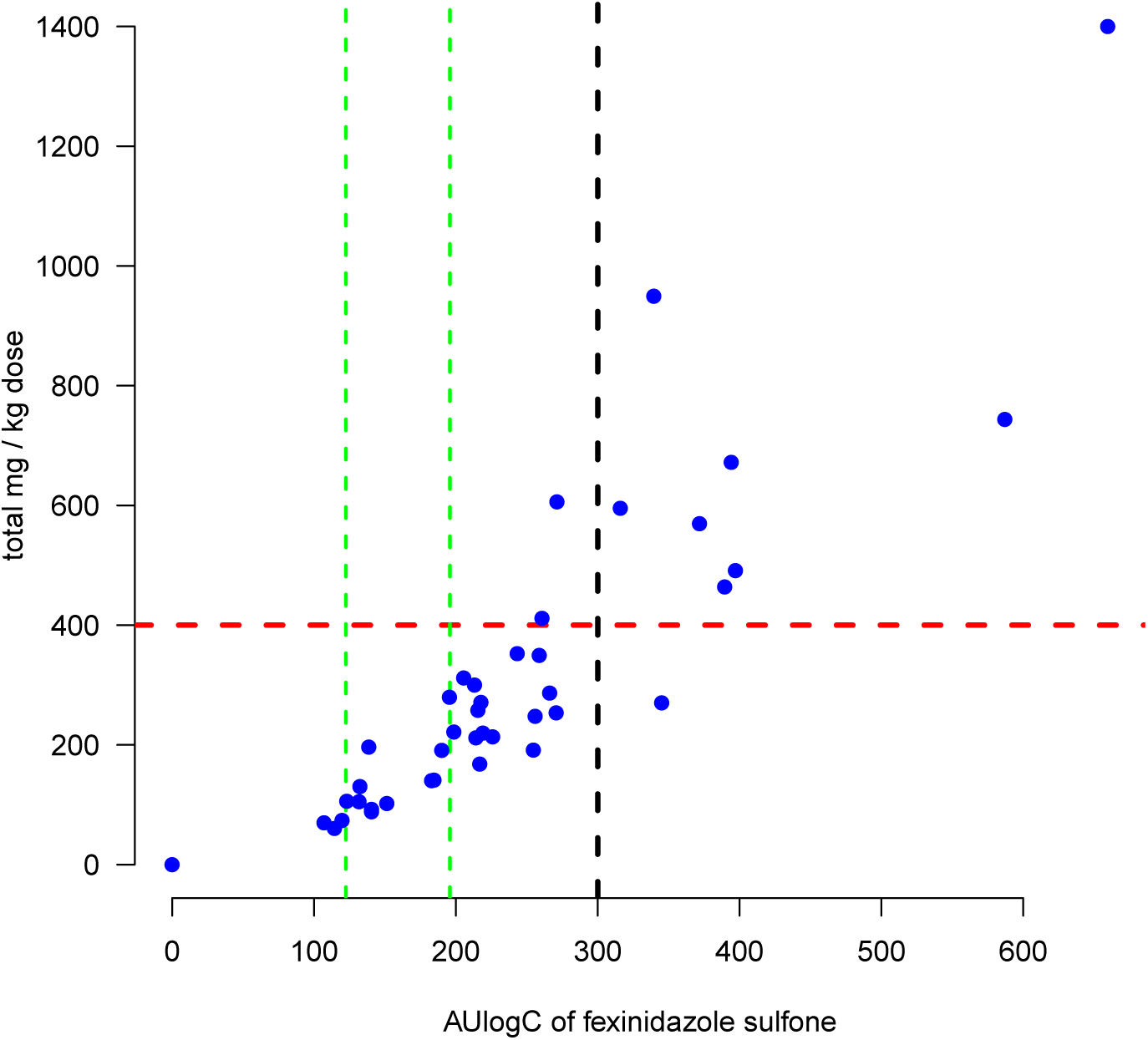
Relationship between the mg/kg dose and the total exposure to fexinidazole sulfone in Chagas disease patients. The vertical green lines show the upper and lower ranges of fitted AUlogC exposures from plasma concentration data in *g*-HAT fexinidazole treated patients. The blue dots show the relationship between mg/kg total dose received and the AUlogC exposure. The black vertical line shows the estimated EC50 for the neutropenia drug effect. The horizontal red line shows our proposed safety cutoff threshold.

**S2 Fig.**
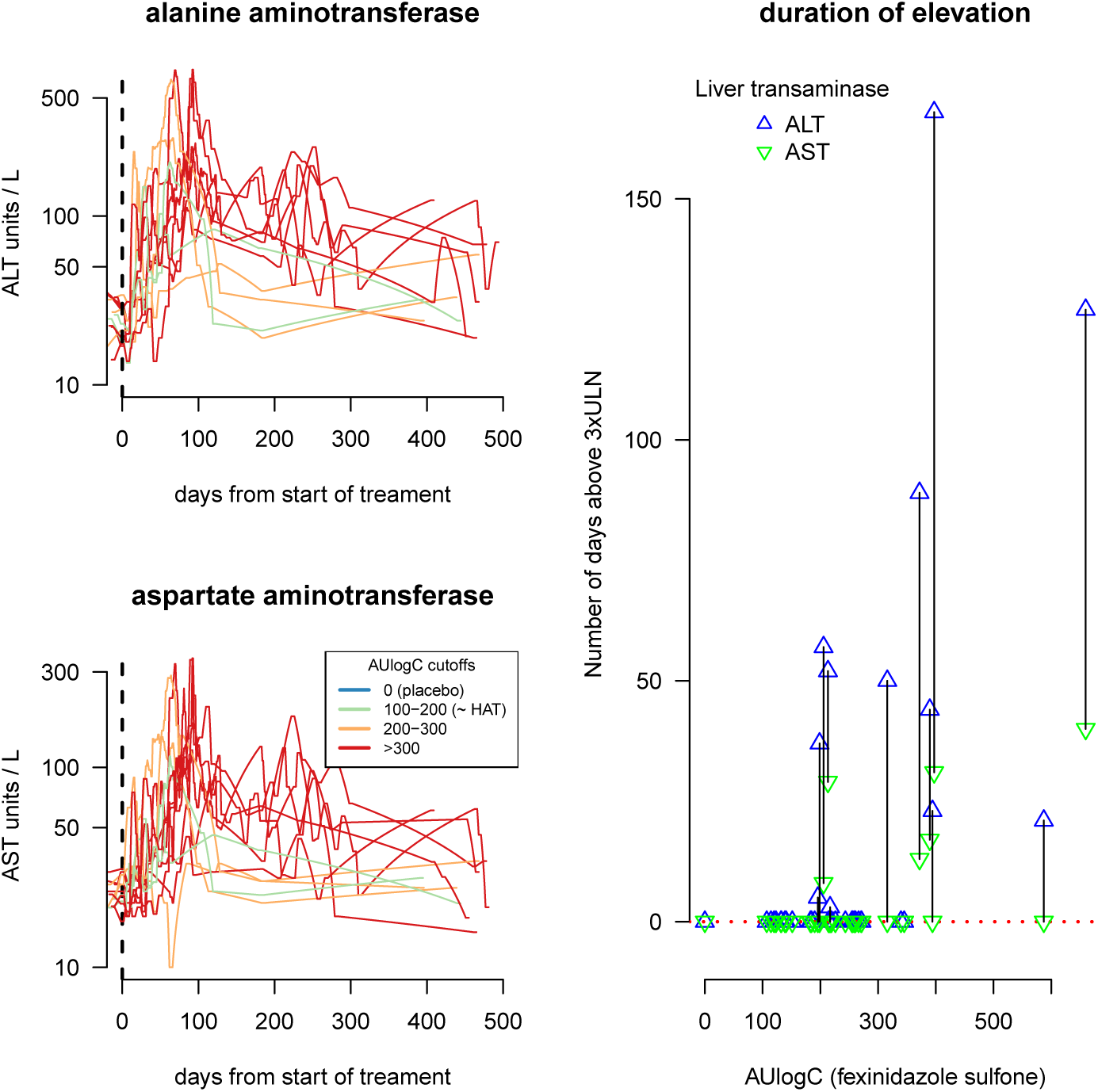
The duration of chronic elevated liver transaminases. The two left panels show the time-series data for ALT (top) and AST (bottom) over the 500 days of follow-up post start of treatment. The right panel shows the relationship between pharmacokinetic exposure and duration of chronic elevation defined as the number of days above 3 x upper limit of normal (ULN). Black lines connect distinct durations of elevations for AST and ALT within the same individuals.

**S3 Fig.**
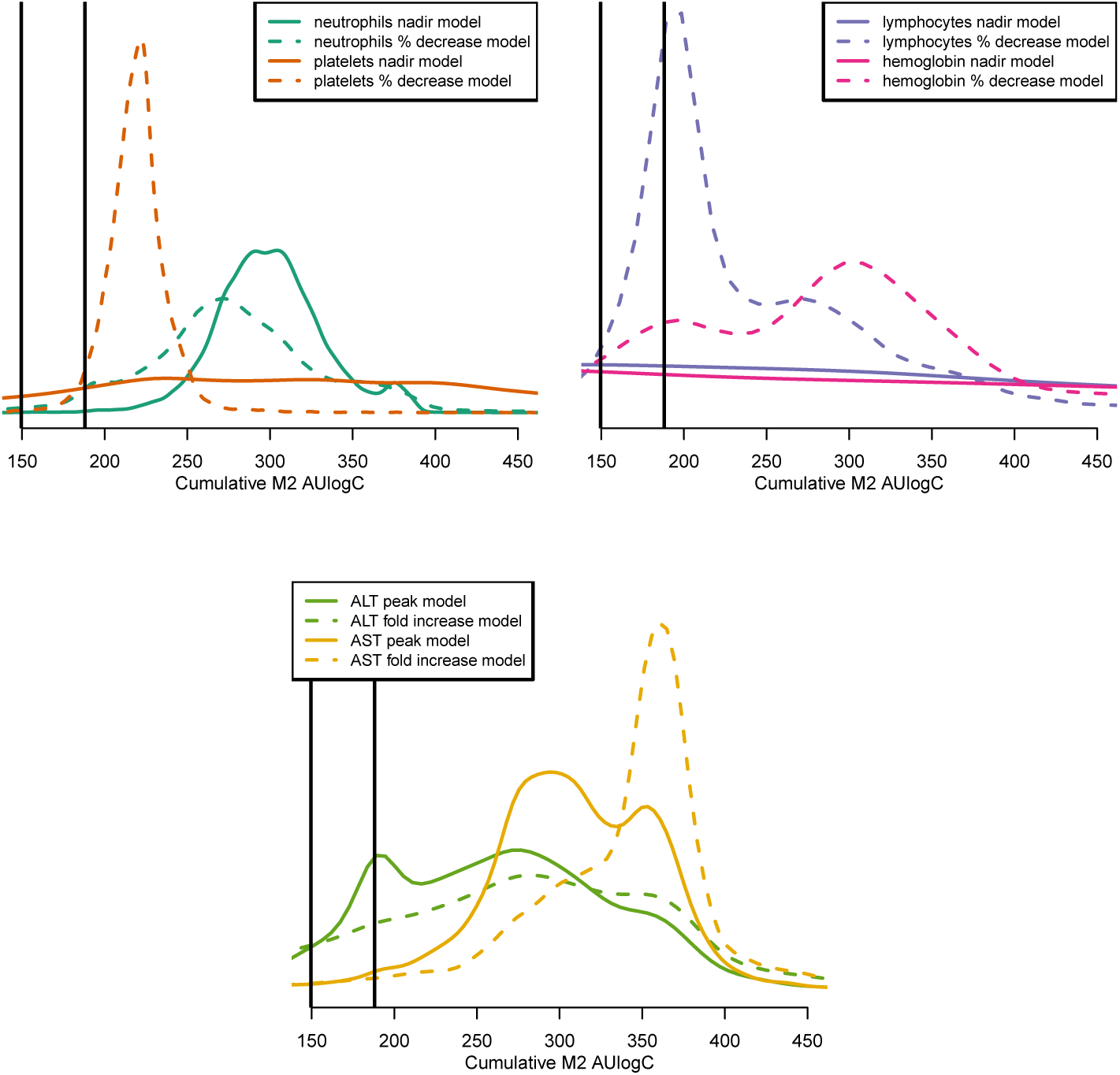
Posterior distribution of *ED*_50_ values for the fitted PK-PD relationships. The vertical black lines show the ranges of exposures observed in the HAT regimen. The colored lines show the posterior distributions for each model. Dashed lines show the posterior fits for the *relative* models and the thick lines for the *absolute* models.

**S4 Fig.**
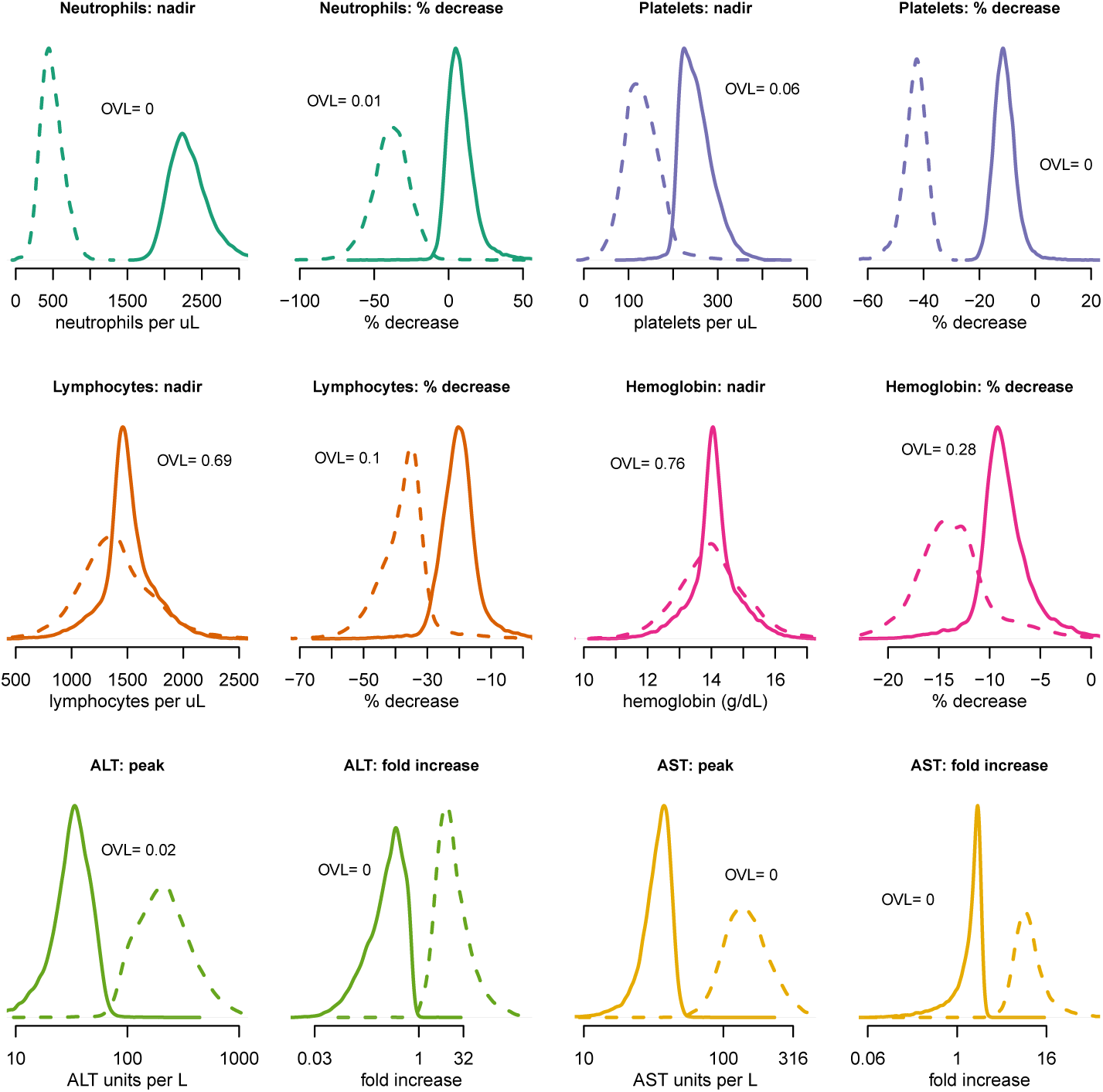
Posterior distributions over the *E*_min_ (thick lines) and *E*_max_ (dashed lines) values for all models and for all pharmacodynamic outcomes of interest. The overlap coefficients (see equation 2) are given in text for each pair of marginal posterior distributions.

**S5 Code NONMEM code used to the fit the final structural model to all M2 data**.

**S6 Code Stan model code used to fit the exposure response relationships.** This gives the full prior distributions used to fit these PK-PD models.

## Acknowledgments

This work was done as part of the Mahidol Oxford Tropical Medicine Research Programme funded by the Wellcome Trust. We are very grateful to Richard Hoglund for his guidance on the pharmacokinetic modeling.

We thank all those involved in the clinical trials of fexinidazole for Chagas and HAT, particularly the patients and healthy volunteers.

The Drugs for Neglected Diseases *initiative* (DND*i*) is grateful to its donors, public and private, who have provided funding to DNDi since its inception in 2003. A full list of DND*i*’s donors can be found at http://www.dndi.org/donors/donors/.

We thank Faustino Torrico, Joaquim Gascón, Lourdes Ortiz, Jimy Pinto, Gimena Rojas, Alejandro Palacios, Fabiana Barreira, Bethania Blum, Graeme Bilbe, Alejandro Gabriel Schijman, Michel Vaillant, on behalf of the FEXI-CHAGAS study group.

We thank the Platform for a Comprehensive Care of Patients with Chagas disease in Bolivia, a collaborative project between CEADES of Health and Environment; Universidad Mayor de San Simon in Cochabamba, Bolivia; Juan Misael University Saracho, Tarija, Bolivia; and ISGLOBAL (Barcelona Institute for Global Health, Spain).

We are grateful for the time spent by the Data Safety Monitoring Board: Drs. Oscar Noya (Chairperson), Rodolfo Viotti, Dominique Larrey, Jean-Louis Paillasseur, and Pascal Voiriot. We also thank Frank Andersohn for expert review of the hematological toxicity findings and Paul Watkins for review of liver safety findings.

## Abbreviations

ALT: alanine aminotransferase
AST: aspartate aminotransferase
AUlogC: area under the log plasma concentration time curve
CNS: central nervous system
CSF: cerebrospinal fluid
DBS: dry blood spot
*g*-HAT: *Trypanosoma brucei gambiense* human African trypanosomiasis
M2: fexinidazole sulfone metabolite
NECT: Nifurtimox-Eflornithine combination therapy
NHV: normal healthy volunteers
OVL: overlapping coefficient
PK-PD: pharmacokinetics-pharmacodynamics

